# Targeting therapy-induced senescence across multiple breast cancer subtypes in a metastatic bone-like microenvironment

**DOI:** 10.64898/2026.05.12.724653

**Authors:** Eleane C.B. Hamburger, Saber Ghazizadeh, Federico Cardahi, Jean A. Ouellet, Michael H. Weber, Livia Garzia, Lisbet Haglund, Derek H. Rosenzweig

## Abstract

Chemotherapeutic treatment of breast cancer with Doxorubicin (DOX) can induce tumor and stromal cell senescence leading to therapy-resistance. Senescence-associated secretory phenotype (SASP) promotes secretion of pro-inflammatory and tumorigenic factors causing systemic inflammation. Combined, this can result in immune suppression, tumor growth and secondary spread of cancer. Targeting and removing senescent and cancerous cells using a combination of chemotherapeutic and senolytic drugs may reduce systemic inflammation, improve therapeutic efficacy, and prevent metastasis. Exposure of triple-negative breast cancer (MDA-MB-231), hormone-responsive (MCF-7) and HER2+ (MDA-MB-453) cells, and primary spine osteoblasts to DOX showed significant induction of p21-positive senescent cells. DOX and senolytics (RG-7112, o-Vanillin) treatment of co-culture spheroids showed a significant additive effect in reducing tumor sphere viability and growth, indicating reduced metastatic potential. This was correlated with reduced SASP in triple-negative and hormone responsive lines and decreased levels of senescent cells in all subtypes and primary stromal cells, while proliferation was decreased, and apoptosis increased across all breast cancer subtypes. Future chemotherapeutic treatment in breast cancer models may be optimized by adding senolytic drugs to more effectively clear senescent tumor and stromal cells, reducing risk for relapse and metastatic potential, while allowing for tissue regeneration in the bone metastatic environment.

**Graphical Abstract:** 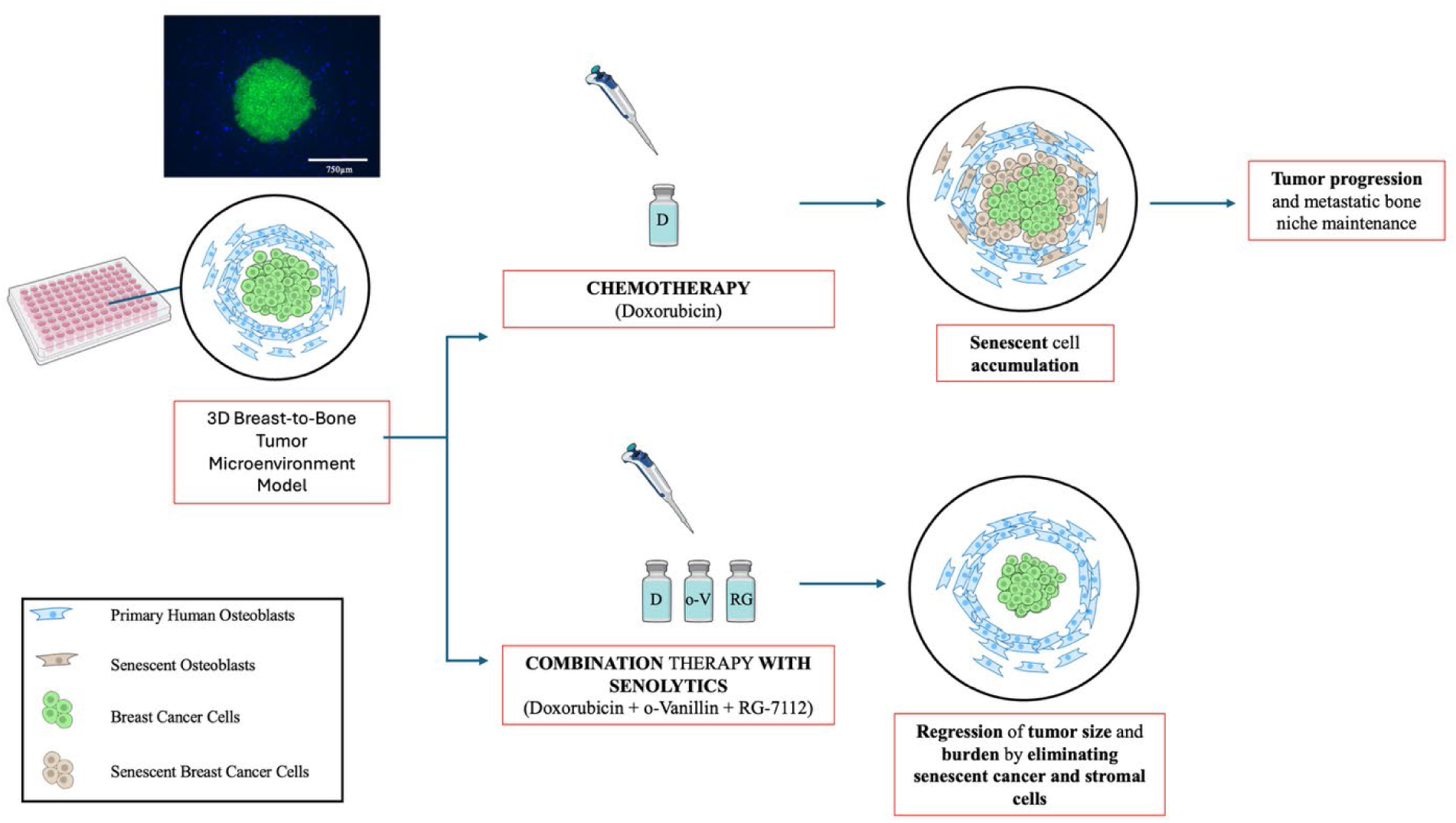

Senolytics selectively eliminate senescent cancer and stromal cells and enhance Doxorubicin efficacy in a 3D bone-like tumor microenvironment model.

## 1. Introduction

Primary breast cancer (BC) is the most frequently diagnosed malignant cancer among women worldwide.[1–3] It commonly metastasizes to bone, most often to the vertebral bodies of the spine, where factors such as receptor activator of nuclear factor κB ligand (RANKL) activate osteoclastic cells, playing a key role in invasion and skeletal events.[4–9] The incidence rate for breast to spine metastasis is 85%, and among the leading causes of metastatic-related death in women.[10] Recent advancements in oncological radiotherapy and surgical treatment have provided a longer life-expectancy for patients post-diagnosis[11], suggesting a more chronic disease. The quality of life for patients with spine metastases is compromised by neurological and structural deficits leading to heightened pain, instability and/or paralysis.[12, 13] The most common subtype of BC is hormone-receptive luminal type A, representing 50-60% of cases, and is defined by estrogen receptor (ER) and/or progesterone receptor (PR) positivity and human epidermal growth factor receptor 2 (HER2) negativity.[14] HER2+ (overexpressing HER2 protein) and triple-negative (lacking ER, PR and HER2 protein expression) BCs account for 10-15% and 15-20% of diagnoses respectively. HER2+ is commonly treated with a combination of monoclonal antibodies such as trastuzumab and chemotherapy. While a majority of patients initially respond well to this treatment, many have continued disease progression within one year, requiring further treatment modalities.[15, 16] The triple-negative subtype lacks the receptors needed for targeted treatment, resulting in a more difficult to treat, and more aggressive type of cancer (40% recurrence)[17], as compared to HER2+ (25% recurrence) and hormone-responsive cancer (5% recurrence rate).[18, 19] Incidence of distant metastatic development for triple-negative BC subtype is in excess of 33%, hormone-responsive 20-30%, and 10-20% in patients with HER2+ BC subtype.[20–22] Many patients that develop breast to spine metastases have been treated with chemotherapy for their primary tumors, possibly driving therapy resistance and maintenance at the metastatic site.

Chemotherapeutic stress has been documented to induce a population of cancer and stromal cell senescence.[23–25] Cellular senescence is a process of stable cell-cycle arrest mediated by the p53-p21-p16 axis.[26, 27] An accumulation of senescent stromal and cancer cells leads to the adoption of a senescence-associated secretory phenotype (SASP).[23, 24, 28] SASP release includes pro-inflammatory and pro-tumorigenic factors such as IL-1β, IL-6, IL-8, IFN-γ and TNF-α.[29, 30] The immunosuppressive and pro-tumorigenic effects of the SASP therefore represent a potential therapeutic target. Doxorubicin (DOX) is an anthracycline that blocks DNA topoisomerase II and is used in standard practice for BC treatment. DOX has been noted to cause systemic inflammation and, when used at higher doses, cardiac toxicity.[31] In contrast, use of clinically relevant low-dose DOX avoids cardiac toxicity but can induce a population of senescent cells.[30, 32–34] Low-dose DOX results in lower clearance of cancer cell populations, potentially facilitating treatment resistance and evasion.[35, 36] Senolytic drugs specifically target senescent cells leading to apoptosis, whereas senomorphic agents suppress SASP production.[37] In addition, senolytics could reduce migration and relapse whereby the senescent stroma contributes to a negative feedback loop, through sustained SASP secretion, within the tumor microenvironment.[38] Therefore, both senolytic and senomorphic drugs could prove useful as an adjunct to chemotherapy by eliminating senescent cancer and senescent stromal cell populations while attenuating SASP factor production within the tumor microenvironment.

Our group has previously shown that the combination of senolytic agents RG-7112 (RO5045337) and o-Vanillin can eliminate senescent cells in musculoskeletal tissues, such as intervertebral discs, resulting in reductions in disc degeneration, pain-associated signatures, and inflammatory SASP signaling.[39–43] RG-7112, an inhibitor of the p53-MDM2 interaction, was originally designed for treatment of acute myeloid leukemia.[44] The high doses needed for cancer treatment resulted in several adverse events including hematological toxicity[45, 46], yet lower doses (∼5 μM) are senolytic and considered safe.[47] o-Vanillin, a natural compound, has both senomorphic, anti-inflammatory and senolytic effects, and has been shown to have anti-proliferative, -angiogenic and -metastatic properties.[48–52] o-Vanillin when previously used at doses up to 300 mg/kg, did not result in any significant adverse side-effects. Here, we apply these lower, senolytic doses in an attempt to reduce therapy-induced senescence, attenuate SASP signaling and improve DOX efficacy.[51, 53] In a previous study we documented that combining these senolytics with DOX could suppress tumor sphere growth of triple-negative BC, hormone-responsive BC and HER2+ BC cell lines.[34] Whether this combination of therapeutics retains efficacy when tumor spheres are placed in a more physiological bone/spine metastatic microenvironment and whether the SASP plays a key role remains unclear. This study aims to assess whether combining DOX with senolytic drugs reduces DOX-induced senescence and enhances overall therapeutic efficacy in a 3D breast-to-bone microenvironment model using three subtypes of BC. Of note, BC subtypes are treated with distinct standard therapeutic strategies; therefore, responses to novel combination therapies may vary. We hypothesized that o-Vanillin and RG-7112 would eliminate senescent stromal and cancer cell populations and attenuate SASP signaling, thereby enabling more effective DOX-mediated suppression of spheroid growth.

## 2. Materials and Methods

### 2.1. Cell isolation and in vitro culture

#### 2.1.1. Tissue collection and bone cells isolation

Primary spine osteoblasts were isolated from human lumbar spine tissue with informed consent in collaboration with Transplant Quebec and approved by the ethical review board at McGill University (Institutional Review Board, IRB #2019-4896). Cortical bone chips were enzymatically digested and cultured to allow osteoblast outgrowth, as previously described in detail.[34, 54] Cells were passaged at 80% confluency and used from passage 1-4. Tissue demographics can be seen in Table 1. Based on data collected at the time of tissue harvest, the organ donors were all considered “healthy donors, pain-free and with no known previous cancer”. Of note, previous cancer diagnosis is an exclusion criterion for Transplant Quebec.

**Table 1.** Characterization of vertebral tissue donor demographics.

| Age | Sex | Cause of Death |
| --- | --- | --- |
| 18 | F | Drowning |
| 33 | F | Trauma |
| 54 | F | Cardiac Arrest |
| 60 | F | Cerebral Hemorrhage |
| 73 | F | Stroke |

#### 2.1.2. Cell culture and seeding

Primary spine osteoblasts were procured as mentioned above and green fluorescent-protein (GFP) expressing epithelial breast adenocarcinoma cell line GFP-MDA-MB-231 were provided by the laboratories of professor M. Park at McGill University, MCF-7 and MDA-MB-453 were commercially sourced (HTB-22 and HTB-131 respectively, purchased from American Type Culture Collection ATCC). All cells were cultured in T75 flasks (Sarstedt, TC Flask T75, Stand, Vent. Cap, Germany) in basal media, high-glucose DMEM (Gibco, Canada) supplemented with 10% FBS (Gibco, Canada) and 1% PS (Gibco, Canada). Cells were placed in incubators 37°C and 5% CO_2_ until 80% confluency was reached for passaging and seeding.

##### a) Standard 2D culture

Chamber slides (Nunc^TM^ Lab-Tek II^TM^ ThermoScientific) were incubated with DMEM full-serum media for 24 hours then seeded with 10,000 MDA-MB-231, MCF-7, or MDA-MB-453 cells per well or 15,000/well primary spine osteoblasts and allowed to adhere overnight at 37°C, 5% CO_2_.

##### b) Spheroid 3D culture/co-culture

Tumor spheroids were generated from MDA-MB-231, MCF-7, or MDA-MB-453 cells using Nunc^TM^ 96-well, Nunclon Delta-Treated, U-Shaped-Bottom Microplate (ThermoFisher Scientific, Toronto, ON, Canada), following a previously established high-throughput spheroid protocol based on MDA-MB-231 Cell Line Spheroid Generation and Characterization protocol for HT Assays (ThermoFisher Scientific). After 24 hours, spheroids were embedded in a type I collagen matrix containing primary human osteoblasts to establish 3D tumor-bone co-cultures as previously described.[55] Cultures were matured under standard conditions, transferred to non-adherent plates (Geiner CELLSTAR®, multi-well culture plate, Sigma-Aldrich, Oakville, ON, Canada), and maintained in low-serum (1% FBS) medium. Treatments were administered between days 7 and 21, with media refreshed every four days. MDA-MB-231 cells were stably labeled with GFP, whereas MCF-7 and MDA-MB-453 were labeled with Calcein-AM for fluorescence-based visualization. 3D cultures were labeled to visualize spheroid morphology and quantify size and invasive outgrowth. Calcein staining was used for structural imaging only and not as a live/dead viability assay.

### 2.2. Therapeutic screening with Doxorubicin, o-Vanillin and RG-7112

2D: Doxorubicin (DOX) hydrochloride (Sigma-Aldrich, #44583) solutions were prepared in PBS to obtain stock solutions. Working solutions were then diluted using low-serum DMEM serving as the vehicle control and added to chambers at final concentrations of 0, 0.1, 0.25 and 0.5 µM DOX. 3D co-culture: DOX was used at a concentration of 0.25µM in monolayer and 0.5µM in 3D culture. o-Vanillin solutions were prepared in PBS at stock concentration 0.02M to achieve a working solution of 100µM in full serum DMEM, and RG-7112 was prepared in PBS as a stock solution of 5mM, then diluted to a working concentration of 5µM in full serum DMEM. Spheroids were treated for 14 days with one of the following: no drugs, DOX, or a combination of DOX, o-Vanillin, and RG-7112, all in low-serum media. In 3D co-culture experiments, drugs were diluted in a similar fashion. For the DOX induction monolayer experiment, a concentration of 0.25 µM DOX was used. In the combination treatment experiment, a concentration of 0.5µM DOX was applied and the co-cultures were treated with either no drugs, DOX, or DOX plus o-Vanillin and RG-7112, all in low-serum media.

### 2.3. Metabolic Activity Measurements

Metabolic activity was assessed with AlamarBlue assay as previously described[40] and as per the manufacturer’s instructions (Invitrogen, #DAL1100). Briefly, at the required time point, media was removed and low-serum media with 10% AlamarBlue was added to each well and incubated at 37°C for 6 hours. Fluorescence exposure was set to Ex560nm/Em590nm and measured using a spectrophotometer (Tecan Infinite T200, Männedorf, Switzerland) using the Magellan software. All metabolic activity is shown as a percentage as compared to the experiments control. AlamarBlue assays were performed for the co-culture spheroids on days 3, 7, 10, and 14 of treatment where day 14 was shown with correlated area data at the same timepoint.

### 2.4. Immunohistological analysis

To determine *in vitro* efficacy of drug combinations on spheroids in co-culture experiments, we prepared the spheroids for sectioning and staining. 3D spheroid co-cultures were washed with PBS and then fixed with 4% paraformaldehyde (Sigma-Aldrich, Oakville, ON, Canada), cryoprotected in 10-30% sucrose, and embedded in Tissue-Plus^TM^ optimal cutting temperature compound (OCT) (Fisher Scientific, Canada). They were then flash-frozen in −80°C for cryopreservation. Sections were made at 16-µm-thick slices and mounted on Fisherbrand Superfrost^TM^ Plus slides (Fisher Scientific, Canada) then placed in −20°C. Cryosectioning was performed on a Leica CM1950 Cryostat (Leica Microsystems, Richmond Hill, ON, Canada). Slides were placed on a 50°C heater for 30 minutes and then washed with PBS mixture with 0.1% Triton X-100 (Sigma-Aldrich, Canada) and 1% Tween (Sigma-Aldrich, Canada). Slides were then saturated in a blocking buffer made with a PBS-Tween-Triton mixture, containing 1% bovine serum albumin (Sigma-Aldrich, Canada) and 1% goat serum (depending on the host of the secondary antibody) for 1 hour at room temperature. For 2D, immunopositivity was done using a fluorescence antibody p21 (ab220206, Abcam, Cambridge, MA, USA) and then the slides were mounted with coverslips using Fluoroshield^TM^ with DAPI (F6057-20ML, Sigma-Aldrich, St. Louis, MO, USA) with the same steps as mentioned above removing the step of exposure to heat. 3D cultures slides were incubated at 4°C overnight with the fluorescence antibody p21 (ab220206, Abcam, Cambridge, MA, USA), or p53 (ab1101, Abcam, Cambridge, MA, USA) for senescence quantification and visualized using a fluorescent secondary antibody Alexa Fluor^TM^ 555 anti-mouse (Invitrogen, ThermoFisher Scientific Waltham, MA, USA) and were then mounted with coverslips using Fluoroshield^TM^ with DAPI (F6057-20ML, Sigma-Aldrich, St. Louis, MO, USA). For proliferation and apoptotic analysis, samples were incubated with primary Ki-67 rabbit mAb (ABclonal Technology, Woburn, MA, USA) or Caspase-3 (ABclonal Technology, Woburn, MA, USA) respectively, at 4°C overnight and then Alexa Fluor^TM^ 555 anti-rabbit (Invitrogen, ThermoFisher Scientific Waltham, MA, USA) conjugated secondary antibody for 2 hours at room temperature. Images were captured using an Invitrogen^TM^ EVOS^TM^ M5000 Imaging System (Invitrogen, #AMF5000) for both brightfield and fluorescent photomicrographs.

### 2.5. Tumor spheroid image analysis

Area of spheroids were measured using Fiji ImageJ version 1.0 (NIH, Maryland, USA) using ROI manager spheroid drawn area to measure and track spheroid growth over time. In the co-culture experiments, we applied a TopHat Fast Fourier Transform (FFT) filter (2.0 pixels) then inversed the FFT of the model image and measured fluorescence intensity and area under the curve from their corresponding 3D surface plot and ROI manager multi plot. Fluorescence intensity and area were measured at 3 days and 14 days post treatment.

### 2.6. SASP factor analysis

Fourteen inflammatory mediators of interest were analyzed using custom Human ProcartaPlex Mix&Match kit (#PPX-15-MXMFZR7) (ThermoFisher Scientific, Toronto, ON, Canada) and measured with the Luminex xMAP system (Serial #MAGPX11188004) (ThermoFisher Scientific, Toronto, ON, Canada) according to instructions given by the manufacturer as previously described.[56] Samples were pre-diluted 1:2 with DMEM media. Standard curves for each analyte were generated with a 4-parameter logistic (4-PL) algorithm. Both median fluorescence intensity (MFI) corrected with background, and final concentrations (pg/mL) were reported by the ThermoFisher ProcartaPlex analysis platform. Few concentrations were out of the detection range and were extrapolated using the associated standard curves for further analysis.

### 2.7. Statistical Analysis

Data were analyzed using GraphPad Prism version 10 (GraphPad, La Jolla, CA, USA). Paired or unpaired t-tests were used when comparing two groups, and multiple pairwise comparisons (one-or two-way ANOVA) were run to evaluate multiple groups. All experiments were run four-five independent times with different osteoblast donors for per ‘*n*’ done in duplicate or triplicate. Statistical analysis was done with the *p*-value set to < 0.05 presenting data as +/- SEM.

## 3. Results

### 3.1. Impact of senolytics on Doxorubicin treatment of senescent breast cancer cells

We previously demonstrated that 72 h exposure to 0.25 µM DOX could simulate therapy-induced senescence in primary human osteoblast and triple negative breast cancer cells by about 2.5-fold and 5-fold respectively [57]. To determine whether senolytics could reduce senescent burden in multiple breast cancer subtypes, 3 different BC cell lines and primary osteoblasts were DOX-induced for 72 h in the presence or absence of o-Vanillin, RG7112 or their combination and p21 expression was assessed as a marker of senescence (Figure 1). Triple-negative MDA-MB-231 exposure to DOX resulted in 14.7 ± 0.51% p21 positive cells; DOX combined with RG-7112, o-Vanillin or a combination of all three reduced the levels to 6.34 ± 0.62%, 8.60 ± 0.39%, and 1.59 ± 0.54% p21 positive cells, respectively. Hormone responsive MCF-7 cell exposure to DOX resulted in 14.32 ± 1.17% p21 positive cells; DOX combined with RG-7112, o-Vanillin or a combination of all three reduced the levels to 7.26 ± 0.48%, 9.14 ± 0.81%, and 2.06 ± 0.17% p21 positive cells, respectively. HER2-positive MDA-MB-453 cell exposure to DOX alone resulted in 16.50 ± 1.85% p21 positive cells; DOX combined with RG-7112, o-Vanillin or a combination of all three reduced the levels to 6.36 ± 0.35%, 7.14 ± 0.45%, and 2.34 ± 0.17% p21 positive cells, respectively. Figure S1 shows the representative immunofluorescent images from which data were quantified. The greatest reduction in p21 positivity in all three BC cell types was observed with the triple-combined treatment of DOX, RG7112 and o-Vanillin. The results were statistically significant, as indicated in Figure 1. Exposure of primary human spine osteoblasts to DOX also significantly increased p21 positivity to 26.74 ± 2.51%, DOX combined with RG-7112, o-Vanillin or a combination of all three reduced the levels to 13.51 ± 1.44%, 21.24 ± 1.78%, and 6.51 ± 0.72% of total cells, respectively.

**Figure 1.**
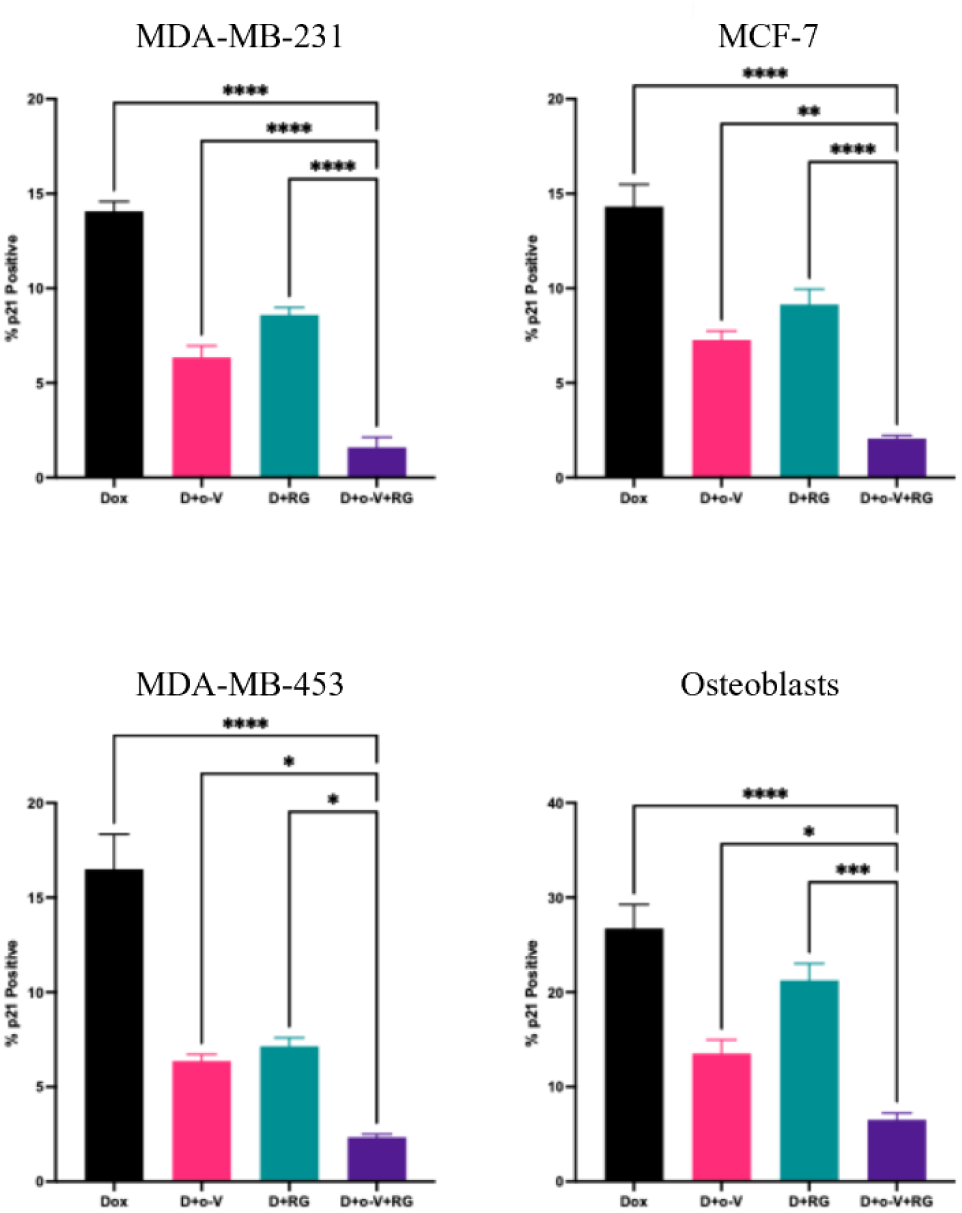
DOX induction and senolytic blockage of senescence in MDA-MB-231, MCF-7, MDA-MB-453 and primary spine osteoblasts. 2D cultures were treated with 0.25 μM Doxorubicin, 5 μM RG-7112, 100 μM o-Vanillin alone or combined for 72h. Graphs indicate quantification of positive p21 stained cells in 2D. n = 4, Mean +/- SEM, * = p < 0.05, ** = p < 0.005, *** = p < 0.001, **** = p < 0.0001, One-way ANOVA.

### 3.2. Senolytics disrupt tumor sphere growth in a bone metastatic model

To generate a more physiologically relevant BC bone metastatic microenvironment, BC tumor spheres were suspended in 3D collagen gels seeded with primary spine osteoblasts. We previously showed that increased DOX (0.5 μM) was required to induce senescence in 3D culture [34]. Co-cultures were treated with DOX alone or in combination with senolytics for 14 days. Figure 2A shows representative images of triple-negative MDA-MB-231, hormone-responsive MCF-7 and HER2+ MDA-MB-453 BC subtype co-cultures on day 14. Quantification of sphere area showed DOX-treated samples were significantly smaller than vehicle-controls for MDA-MB-231 and MCF-7 co-cultures 33606 ± 1334.15 pixels^2^ and 4580.5 ± 884.32 pixels^2^ versus controls 81635.5 ± 10318.15 pixels^2^ and 8552.5 ± 1496.95 pixels^2^. There was further reduction of area across all subtypes with the combination 25869 ± 2726.88 pixels^2^, 1930 ± 151.76 pixels^2^, and 1384.63 ± 406 pixels^2^ (Figure 2B). As a ratio to control, exposure to DOX alone or the triple combined treatments resulted in a significant reduction of sphere area of 0.43 ± 0.04 units (2.33-fold) and 0.33 ± 0.05 units (3.03-fold), for triple negative BC, a decrease of 0.81 ± 0.08 units (1.23-fold) and 0.57 ± 0.04 units (1.75-fold), for hormone responsive BC and no decreases for the HER2+ BC, 1.23 ± 0.04 units and 1.06 ± 0.07 units. After 14 days of treatment, there was also a significant corresponding reduction in metabolic activity compared to vehicle-controls where DOX treated samples showed a decrease of 0.43 ± 0.07 units (2.33-fold), 0.29 ± 0.05 units (3.45-fold), and 0.16 ± 0.02 units (6.25-fold).The triple-combined treatment showed reduction in metabolic activity of 0.03 ± 0.04 units (33.3-fold), 0.03 ± 0.002 units (33.3-fold), and 0.16 ± 0.06 units (6.25-fold) in triple negative, hormone responsive and HER2+ BCs respectively(Figure 2C). These data not only show the expected impact of DOX on BC sphere growth and metabolic activity but strikingly show an additive benefit of combing o-Vanillin and RG-7112.

**Figure 2.**
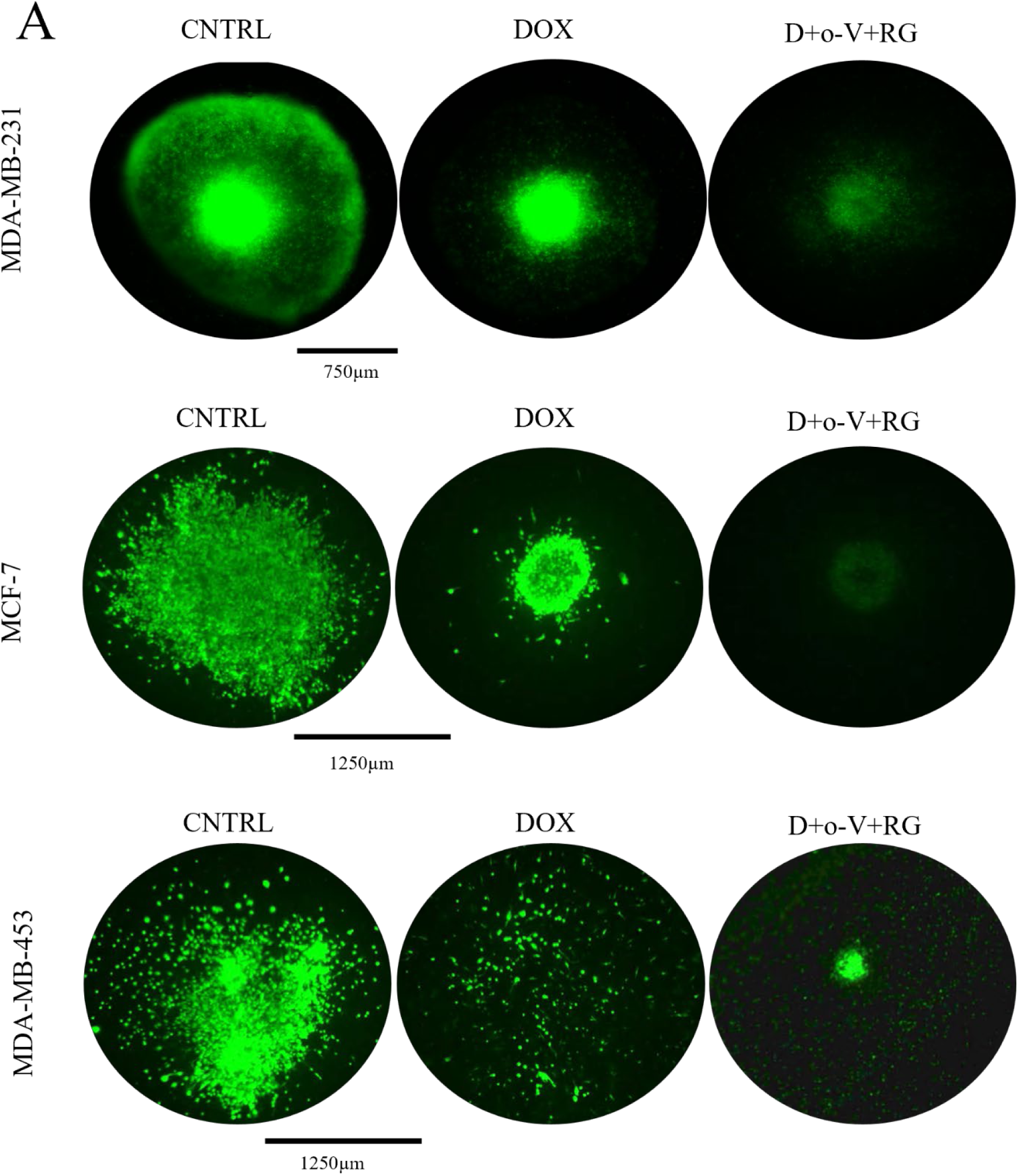
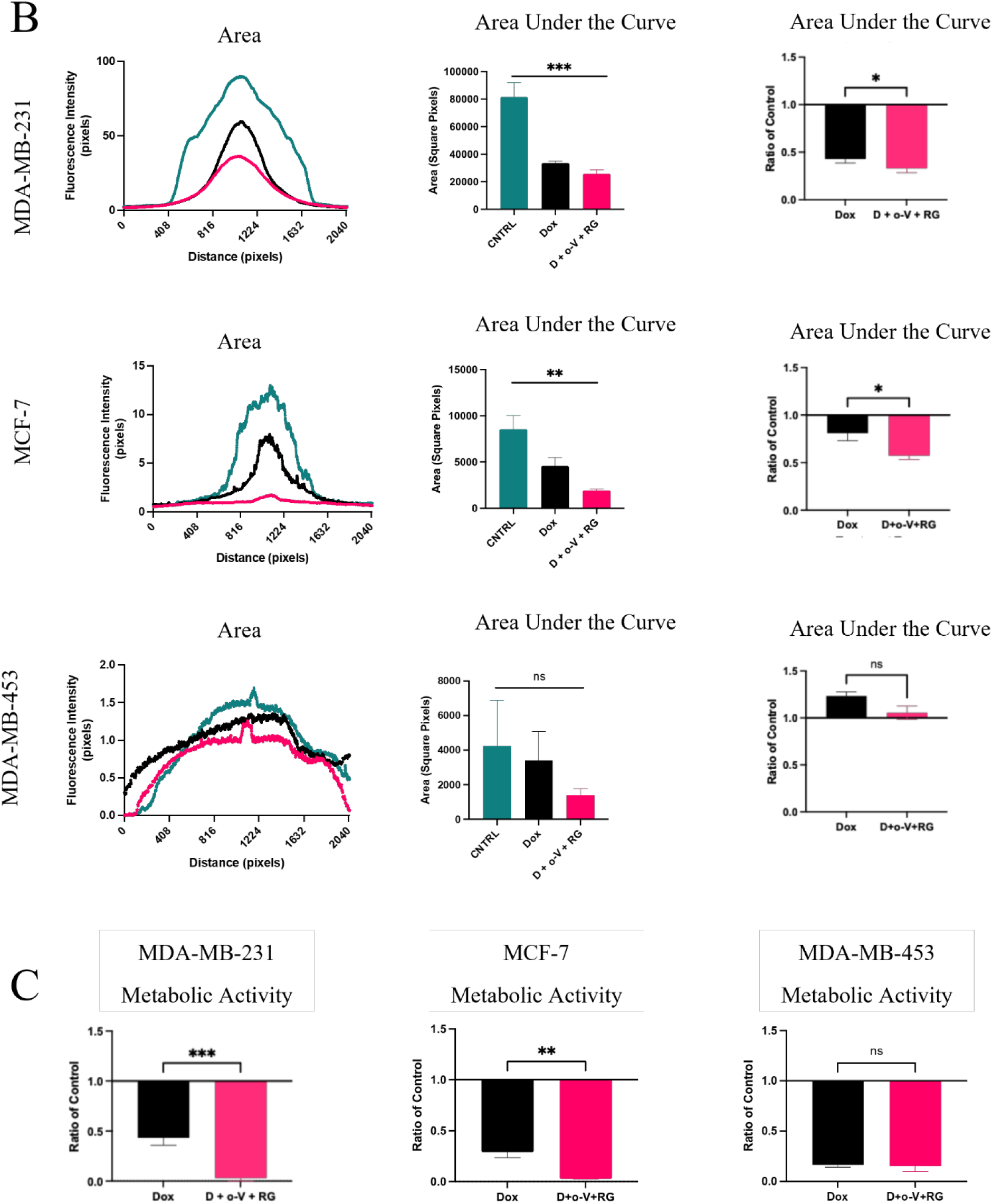
Tumor spheroid growth and metabolic activity in a bone metastatic microenvironment treated 14 days with Doxorubicin alone or in combination with o-Vanillin and RG-7112. A) Representative photomicrographs of GFP-tagged MDA-MB-231 spheroids and Calcceiein-AM MCF-7 and MDA-MB-453 spheroids after 14 days of treatment. B) Tumor spheroid area with quantification of area under the curve. C) Tumor spheroid metabolic activity. n = 4, Mean +/- SEM. * = p < 0.05, ** = p < 0.005, *** = p < 0.001, **** = p < 0.0001, one-way ANOVA with Tukey, unpaired t-test.

To determine whether the striking impact of adding senolytics was additive or synergistic to DOX, the triple-combined treated co-culture sphere size and metabolic activity were compared to the DOX-alone treated samples (Figure 3). The combined treatment showed significant synergistic decrease in metabolic activity across the three BC subtypes by 0.12 ± 0.03 units (8.33-fold) for triple-negative BC, 0.10 ± 0.02 units (10-fold) for hormone responsive BC and no impact on metabolic activity of 0.93 ± 0.27 units (1.07-fold) for the HER2+ BC compared to DOX-alone treated samples (Figure 3). These data indicate that combining senolytics o-Vanillin and RG-7112 with DOX may have a significant synergistic effect on disrupting triple negative and hormone responsive breast cancer growth and activity in a bone-like microenvironment.

**Figure 3.**
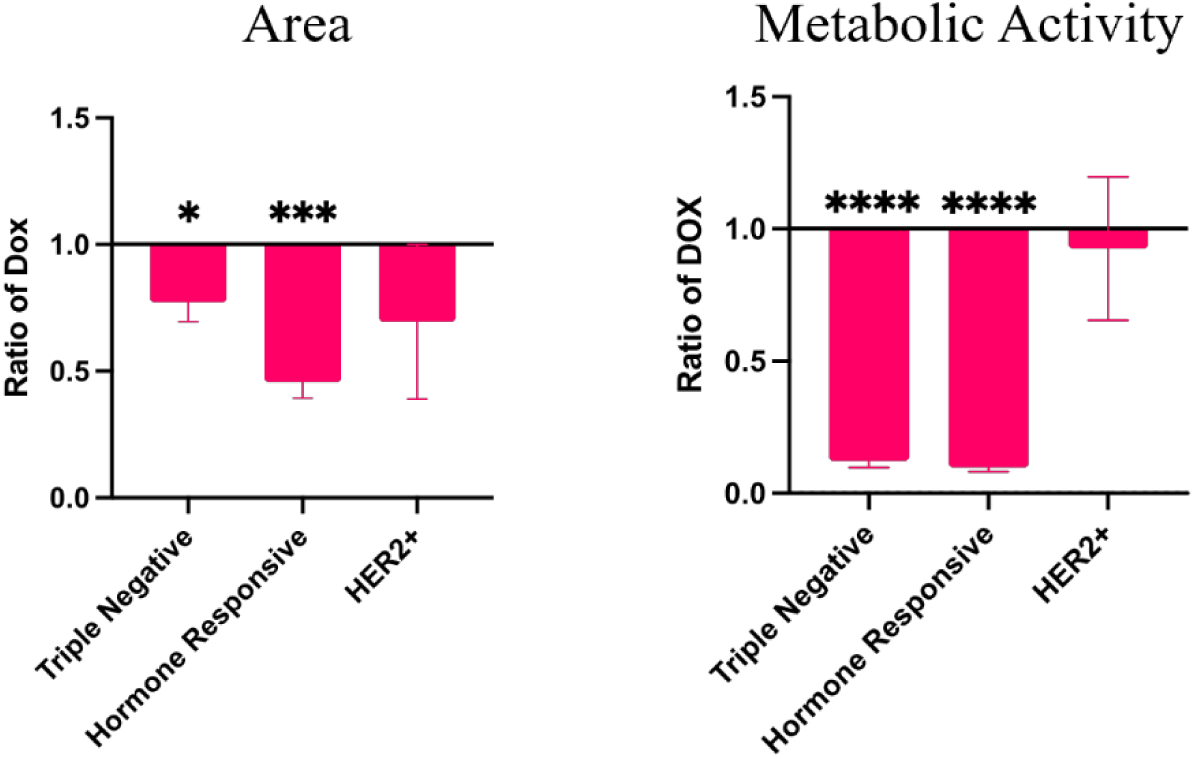
Cross comparison of the impact of combined treatment versus doxorubicin on spheroid growth and metabolic activity in a bone-like microenvironment. Osteoblast cocultures with the three BC subtypes were treated 14 days with combination of DOX with o-Vanillin and RG-7112. Tests for synergy were performed. n = 4, Mean +/- SEM. * = p < 0.05, *** = p < 0.001, **** = p < 0.0001, one-way ANOVA with Tukey.

### 3.3. Quantification of DOX-induced senescence and senolytic removal of senescence in the 3D BC spheroid co-cultures

To evaluate if the reduced sphere size and metabolic activity were linked to the removal of senescent cells, in the indicated co-cultured models, we exposed them to DOX and senolytics as above and the senescent markers p21 and p53 were evaluated (Figure 4). Exposure to DOX significantly increased p21 positivity over untreated controls in all cell types to 8.39 ± 1.86% (triple-negtive), 13.52 ± 4.17% (hormone responsive), 22.28 ± 1.96% (HER2+), and 12.95 ± 1.64% (primary osteoblasts) of total cells. For p53, all except hormone responsive cells showed a significant increase of positively expressing cells with DOX treatment over untreated controls at 28.58 ± 5.64% (triple-negative), 20.24 ± 2.58% (hormone responsive), 19.34 ± 2.55% (HER2+), and 6.89 ± 2.02% (osteoblasts) of total cells. The combination treatment significantly reduced the p21 levels to 0.99 ± 0.40% (triple-negative), 4.04 ± 0.66% (hormone responsive), 1.00 ± 0.75% (HER2+), and 3.91 ± 1.72% (osteoblasts) of total cells. The combination treatment significantly reduced the p53 levels to 7.20 ± 3.45% (triple-negative), 5.06 ± 2.44% (hormone responsive), 0.77 ± 0.51% (HER2+), and 0.87 ± 0.49% (osteoblasts) of total cells. Representative images for the p21 and p53 immunofluorescence can be viewed in supplemental Figure S2. These data show that in a more complex 3D tumor-like microenvironment, DOX induces senescence in both the tumor and stromal compartments. Moreover, the addition of the two senolytics brought the senescence levels approximately back to baseline.

**Figure 4.**
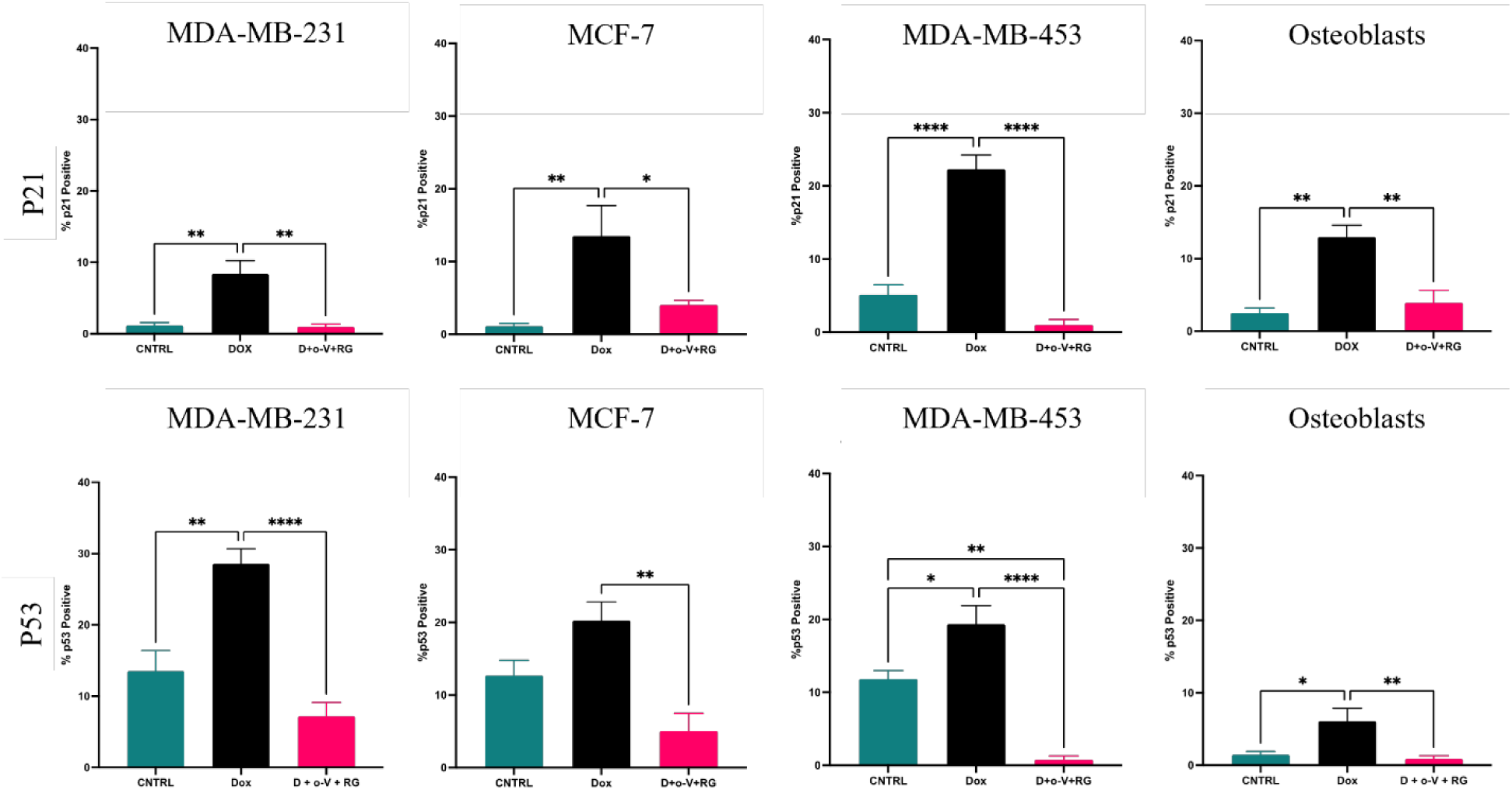
Combination treatment mitigates DOX-induced senescence in BC and primary osteoblast cells. Senescent activity evaluated by p21 and p53 immunofluorescence staining after 14 days of treatment in coculture. Doxorubicin alone significantly increases p21 and p53 levels in all cell types, while the addition of the senolytics significantly reduces the number of p21 and p53 positive cells to basal levels. n = 5, Mean +/- SEM, * = p < 0.05, ** = p < 0.005, *** = p <0.001, **** = p < 0.0001, One-way ANOVA with Tukey.

**Figure 5.**
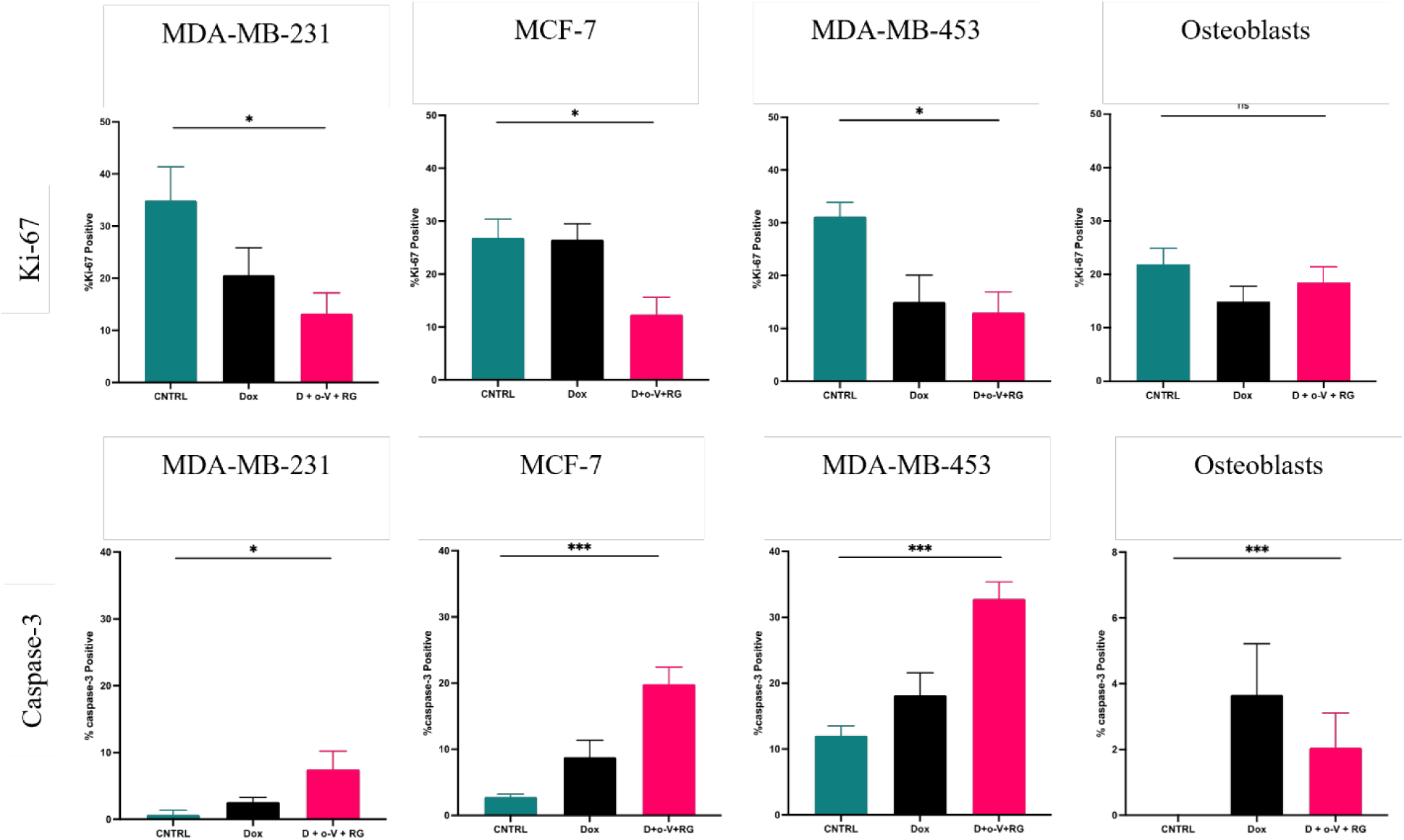
Impact of senolytic treatment of BC spheroid cocultures on proliferation and apoptosis. Proliferation and apoptotic activity evaluated by Ki-67 and Caspase-3 immunofluorescence staining. Green bars indicate untreated controls, black bars indicated DOX alone, and red bars indicate the combined treatment, all over 14days. Osteoblasts were co-cultured with each BC subtype, and assessed in the periphery. n = 5, Mean +/- SEM, * = p < 0.05, ** = p < 0.005, *** = p <0.001, **** = p < 0.0001, Ordinary One-way ANOVA.

**Figure 6.**
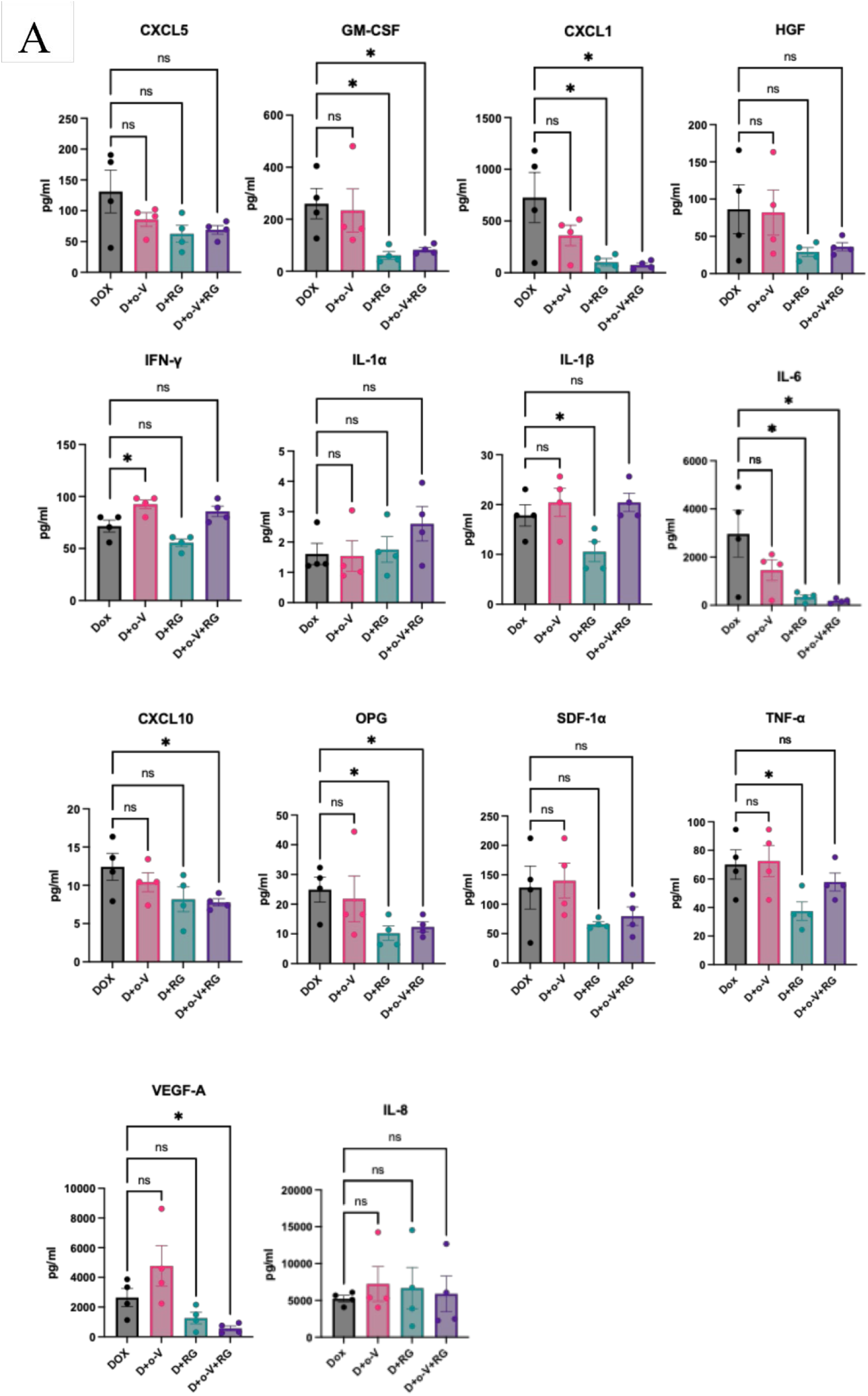
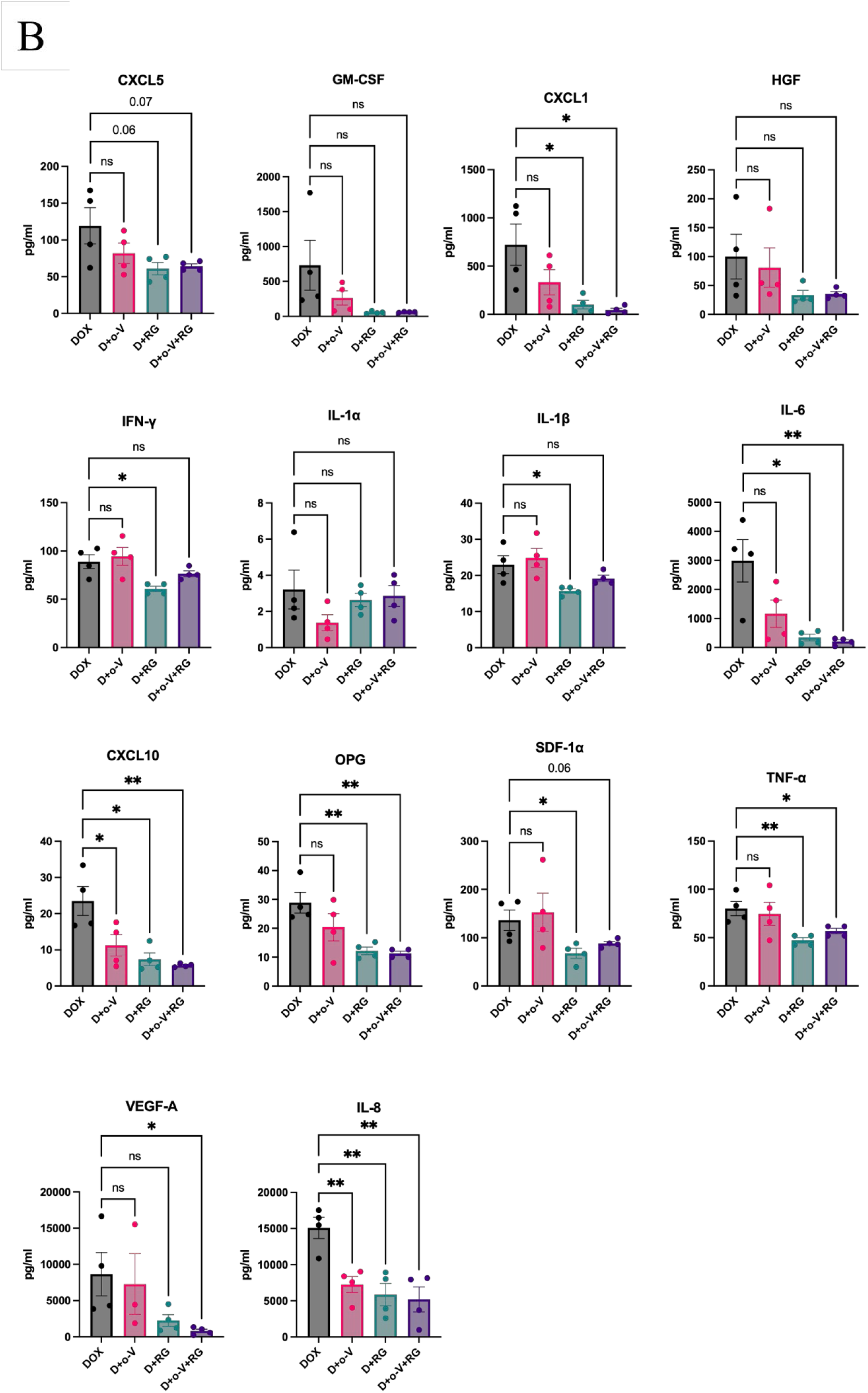
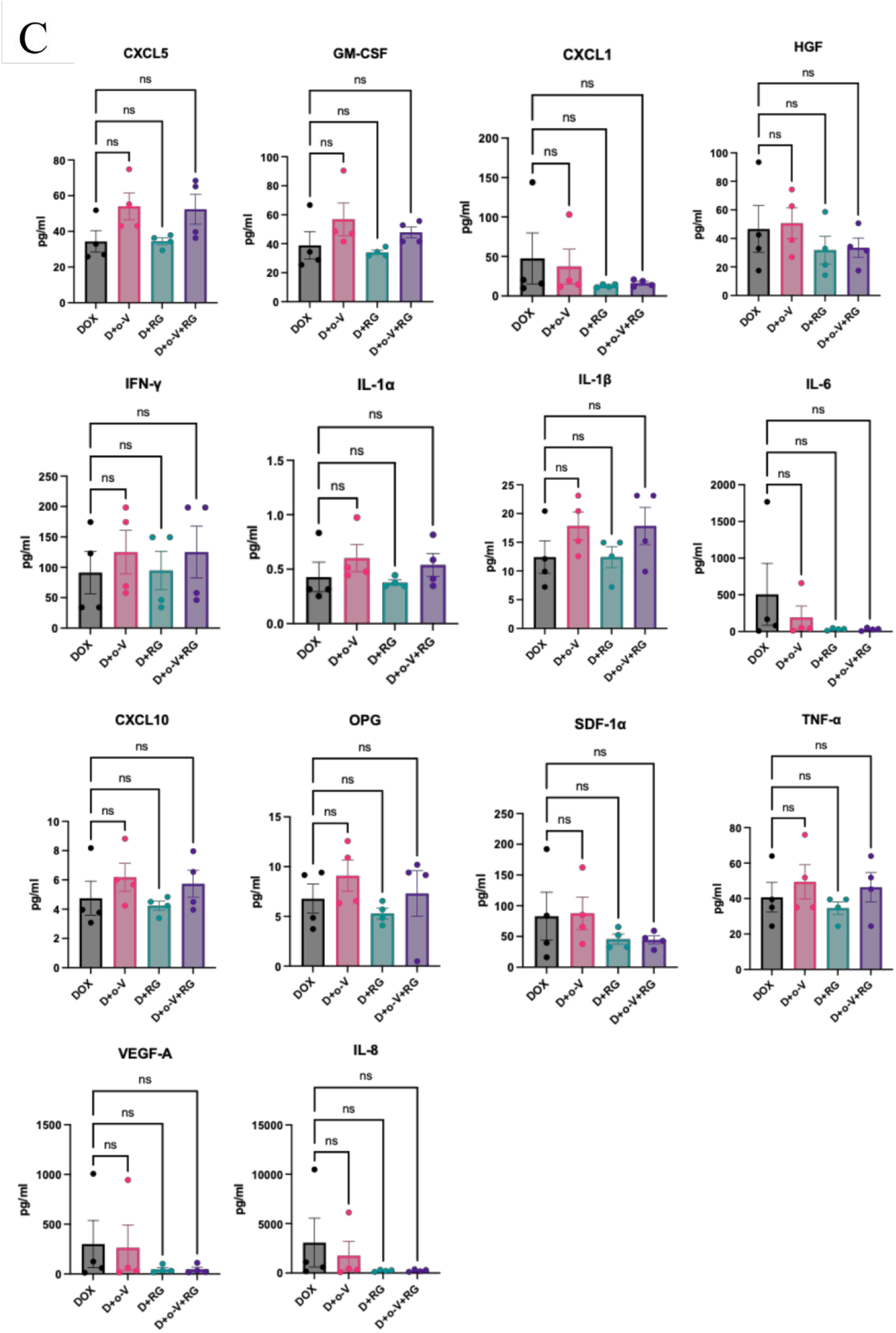
SASP profile analysis of treated BC spheroid/osteoblast co-cultures. Luminex assay for CXCL5, GM-CSF, CXCL1, HGF, IFN-γ, IL-1α, IL-1β, IL-6, CXCL10, OPG, SDF-1α, TNF-α, VEGF-A and IL-8 at 7 days post-treatment. A) MDA-MB-231, B) MCF-7, C) MDA-MB-453. n=4, Mean +/- SEM. * = p < 0.05. For the statistical analysis using GraphPad Prism, unpaired t-test was applied.

### 3.4. Quantification of proliferation and apoptosis following DOX and combination treatments to the BC spheroids in 3D co-culture

To determine whether DOX or combination treatments impact proliferation or apoptosis of the BC cell or primary osteoblasts, positively expressing Ki-67 (proliferation) and Caspase-3 (apoptosis) cells were quantified via immunofluorescence. In the untreated controls, the percentage of proliferation for triple-negative BC was 34.91 ± 6.48%, which was significantly reduced to 20.61 ± 5.27% after DOX treatment, and further reduced to 13.20 ± 3.99% after combined treatment. The percent proliferating hormone responsive BC for untreated controls was 26.85 ± 3.56%, which was similar at 26.52 ± 3.01% after DOX treatment, and significantly reduced to 12.36 ± 3.27% after combined treatment. The percent proliferating HER2+ BC was 31.17 ± 2.70% in controls, with significant reduction to 15.01 ± 5.05% following DOX treatment, and 13.03 ± 3.88% following combined treatment. The caspase-3 marker for apoptosis showed the opposite effect. The triple-negative BC control samples had 0.69 ± 0.60% apoptosis, while DOX treatment significantly increased it to 2.59 ± 0.71% apoptosis, and the combined treatment further significantly increased the apoptosis to 7.48 ± 2.74% of total cells. Hormone responsive BC control samples had 2.78 ± 0.42% apoptotic cells, while DOX treatment significantly increased this to 8.79 ± 2.56% apoptosis, and the combined treatment further significantly increased it to 19.84 ± 2.59% apoptosis. The untreated control HER2+ BC samples had 12.05 ± 1.46% apoptosis, while the DOX treated samples significantly increased to 18.17 ± 3.39% apoptosis, and combined treatment further significantly increased this to 32.79 ± 2.57% apoptosis. Osteoblast proliferation was not affected showing 21.87 ± 3.00% in controls, 14.93 ± 2.86% in DOX-treated and 18.50 ± 2.94% in combined treated samples. The corresponding apoptotic quantification for control, DOX-and combined-treatment samples also showed no significant reduction with 0%, 3.66% and 2.05% apoptotic cells, respectively.

### 3.5. Quantification of SASP factor secretion following DOX-induction and combination treatment to BC spheroid co-cultures

To determine the impact of DOX-induction and addition of senolytics on SASP factor secretion from the BC spheroid co-cultures, conditioned medium was collected after 7 days of treatment and assessed by Luminex multiplex assay. Furthermore, since o-Vanillin and RG7112 impact different signaling pathways and may result in altering different cytokine/chemokine release, single arm senolytic treatments were also assessed in addition to the combined treatment. A panel of known SASP factors involved in tumor microenvironment were probed: CXCL5, GM-CSF, CXCL1, HGF, IFN-**γ**, IL-1α, IL-1β, IL-6, CXCL10, OPG, SDF-1α, TNF-α, VEGF-A and IL-8.

For the triple-negative BC co-cultures (Figure, the combination with RG-7112 and the combination treatment with o-Vanillin plus RG-7112 each resulted in a significant reduction in six out of the fourteen factors. The combination with o-Vanillin only showed a significant increase in factor IFN-**γ,** while all other factors showed no significant change. When compared to DOX alone, combination with RG-7112 significantly reduced GM-CSF (260.00 ±_58.14 pg/mL vs 61.41 ± 14.87 pg/mL (p = 0.016)), CXCL1 (726.70 ± 242.90 pg/mL vs 101.80 ± 35.98 pg/mL (p = 0.044)), IL-1β 17.83 ± 2.14 pg/mL vs 10.56 ± 2.00 pg/mL (p = 0.048)), IL6 (2964.00 ± 981.90 pg/mL vs 338.80 ± 100.70 pg/mL (p = 0.038)), OPG (24.88 ± 4.18 pg/mL vs 10.21 ± 2.42 pg/mL (p = 0.023)), TNF-α (70.12 ± 10.26 pg/mL vs 37.49 ± 6.48 pg/mL (p = 0.036)). When compared to DOX alone, combination with o-Vanillin plus RG-7112 significantly reduced GM-CSF (260.00 ± 58.14 pg/mL vs 82.51 ±_8.82 pg/mL (p = 0.023)), CXCL1 (726.70 ± 242.90 pg/mL vs 75.22 ± 19.37 pg/mL (p = 0.037)), IL6 (2964.00 ± 981.90 pg/mL vs 182.30 ± 48.10 pg/mL (p = 0.030)), CXCL10 (12.42 ± 1.77 pg/mL vs 7.765 ± 0.47 pg/mL (p = 0.044)), OPG (24.88 ± 4.18 pg/mL vs 12.36 ± 1.64 pg/mL (p = 0.032)), VEGA (2642.00 ± 608.70 pg/mL vs 570.50 ± 164.00 pg/mL (p = 0.017)). IFN-**γ** was slightly but significantly increased by DOX plus o-Vanillin treatment (71.59 ± 5.73 pg/mL vs 92.53 ± 4.30 pg/mL (p = 0.027)).

For the hormone responsive BC co-cultures, the combination with o-Vanillin resulted in two significant reductions of the fourteen factors. The combination with RG-7112 resulted in a significant reduction in nine of the fourteen factors. Further, the triple-combination treatment resulted in a significant reduction in seven of the fourteen factors. The combinations with RG-7112 and o-Vanillin plus RG-7112 in CXCL5 when compared to DOX, although not significant, showed a clear trend towards reduction (119.10 ± 24.79 pg/mL vs 60.91 ± 8.56 pg/mL (p = 0.068), and vs 64.31 ± 3.12 pg/mL (p = 0.071)). Further, SDF-1α showed a non-significant reduction when DOX was compared to the triple combination (136.40 ± 21.21 pg/mL vs 88.30 ± 4.17 pg/mL (p = 0.068)). When compared to DOX alone, combination with o-Vanillin significantly reduced CXCL10 (23.48 ± 3.99 pg/mL vs 11.24 ± 2.95 pg/mL (p = 0.049)), and IL-8 (15093 ± 1473 pg/mL vs 7263 ± 1116 pg/mL (p = 0.006)). When compared to DOX alone, combination with RG-7112 significantly reduced CXCL1 (722.00 ± 214.30 pg/mL vs 101.10 ± 44.69 pg/mL (p = 0.030)), IFN-**γ (**89.00 ± 7.23 pg/mL vs 60.73 ± 2.88 pg/mL (p = 0.011)), IL-1<u>β</u> (22.98 ± 2.46 pg/mL vs 15.70 ± 0.64 pg/mL (p = 0.029)), IL-6 (2986 ± 732.60 pg/mL vs 341.40 ± 113.4 pg/mL (p = 0.012)), CXCL10 (23.48 ± 3.99 pg/mL vs 7.38 ± 1.77 pg/mL (p = 0.010)), OPG (28.89 ± 3.60 pg/mL vs 12.21 ± 1.33 pg/mL (p = 0.005)), SDF-1α (136.40 ± 21.21 pg/mL vs 67.91 ± 10.42 pg/mL (p = 0.027)), TNF-α (80.04 ± 7.37 pg/mL vs 47.29 ± 2.79 pg/mL (p = 0.006)), and IL-8 (15093 ± 1473 pg/mL vs 5860 ± 1545 pg/mL (p = 0.005)). When compared to DOX alone, combination with o-Vanillin plus RG-7112 significantly reduced CXCL1 (722.00 ± 214.30 pg/mL vs 44.67 ± 19.79 pg/mL (p = 0.020)), IL-6 (2986 ± 732.6 pg/mL vs 207.10 ± 59.67 pg/mL (p = 0.009)), CXCL10 (23.48 ± 3.99 pg/mL vs 5.72 ± 0.24 pg/mL (p = 0.004)), OPG (28.89 ± 3.60 pg/mL vs 11.34 ± 0.76 pg/mL (p = 0.003)), TNF-α (80.04 ± 7.37 pg/mL vs 56.90 ± 2.76 pg/mL (p = 0.026)), VEGF-A (8653.00 ± 2993.00 pg/mL vs 802.30 ± 250.20 pg/mL (p = 0.040)), and IL-8 (15093 ± 1473 pg/mL vs 5189 ± 1738 pg/mL (p = 0.005)).

For the HER2+ BC co-culutres, the combinations resulted in no significant changes when compared to DOX alone for all fourteen SASP factors. Overall, these data indicate that combined treatment reduced about half of all chosen SASP factors in the triple-negative and in hormone receptor-positive co-culture groups, possibly linked to decreased tumor cell growth and proliferative activity. Further, these results suggest differential responses to DOX combination treatment across the three BC subtypes with respect to SASP factor release.

## 4. Discussion and Conclusion

In this study, we tested the hypothesis that senolytics could enhance doxorubicin efficacy by removing DOX-induced senescent tumor and stromal cells and their associated SASP using an *in vitro* 3D BC cancer spheroid-osteoblast co-culture system. Building on our previous study which focused on triple negative BC cells [34], we expanded the present study to include two additional BC subtypes – hormone responsive and HER2 overexpressing cells. We demonstrate that DOX indeed induces senescence in both tumor and osteoblast (stromal) compartments. More importantly, we show that combining o-Vanillin and RG-7112 significantly reduced senescence burden, altered SASP profiles, and suppressed the overall co-culture metabolic activity, thereby enhancing the therapeutic response to DOX. These findings suggest that senolytic combination with chemotherapy could be a potentially improved treatment strategy for patients with breast-to-bone metastases.

As indicated in several other reports [58–63], we found that DOX induced a senescent phenotype across all three BC subtypes. This was characterized by increased p21/p53 expression, reduced proliferation, and growth arrest rather than acute cytotoxicity. Notably, senescence was also induced in the primary human osteoblasts, which were dispersed around the BC spheroids in our co-culture system. This showed novelty in our study as a bone-like microenvironment, extending beyond other studies on chemotherapy-induced senescence that rely on immortalized-fibroblast-dominated *in vitro* tools or animal models.[40, 48, 53, 64, 65] Therapy-induced senescence has been widely reported following chemotherapy and/or radiation across various breast cancer in vitro and in vivo models, yet comparatively few studies have examined senescence induction within the bone metastatic niche.[25, 66, 67] Together with these other studies, our work here reinforce the importance of addressing therapy-induced senescence in a multicellular bone-like microenvironment, especially since patients with bone metastases have often faced both chemotherapy and radiotherapy stressors along their treatment paths. Indeed, both the tumor itself and surrounding stromal compartment actively contribute to senescence persistence, paracrine signaling through SASP, therapy resistance which all lend to lesion site maintenance.

A central objective of this work was to determine whether senolytic intervention could reduce or reshape the SASP output following DOX-induced senescence. Senolytic treatment altered SASP profiles in a subtype-specific manner, with more pronounced suppression of pro-tumorigenic cytokines in triple-negative and hormone receptor-positive co-cultures than in HER2-positive co-cultures. RG-7112 combined with DOX broadly reduced inflammatory SASP components, including IL-1β, IL-6, OPG, TNF-α, and IL-8, consistent with p53-mediated regulation of senescence-associated inflammatory signaling.[68, 69]. In contrast, o-Vanillin exerted more selective effects on factors including CXCL10, IL-8 and IFN-γ in a context-dependent manner, underscoring pathway-specific regulation of SASP output rather than uniform suppression.[70–73] Beyond the factors discussed above, senolytic treatment also modulated additional SASP-associated mediators linked to immune regulation, angiogenesis, and bone remodeling, supporting a broad, but context-dependent attenuation of senescence-associated paracrine signaling. Collectively, these findings highlight that SASP attenuation occurs in a factor-specific and BC subtype-dependent manner, reflecting differential regulation of signaling pathways possibly linked to nuclear factor-κB (NF-κB)-, p53-, and interferon-associated pathways, which may modulate SASP profiles rather than acting as a uniform transcriptional program.[67, 70, 71]

These subtype-specific SASP patterns are consistent with differential engagement of senescence maintenance and clearance pathways, prompting consideration of how o-Vanillin and RG-7112 act through complementary, but non-overlapping mechanisms across genetically distinct BC subtypes. Functionally, reductions in IL-6 and CXCL1 are particularly relevant given their established roles in promoting BC proliferation, migration, and therapy resistance through autocrine and paracrine signaling.[74–76] In particular, IL-6 and IL-8 are established activators of JAK/STAT3 signaling in stromal and immune compartments, contributing to pro-tumorigenic microenvironmental remodeling,[70, 77] while modulation of OPG is mechanistically relevant in bone-associated disease through its interaction with the RANK/RANKL. These factors have been implicated in invasive progression in triple-negative BC.[78] Additional attenuation of TNF-α, VEGF-A, GM-CSF, CXCL10, and OPG suggests that future studies on senolytic interventions should investigate potential involvement of immune dysregulation, angiogenic signaling, and bone-associated tumor-stromal crosstalk using relevant *in vivo* models.[78–81] The observed reduction of these factors coincided with decreased BC spheroid growth and metabolic activity for triple-negative and hormone receptor-positive subtypes, whereas treated HER2-positive co-cultures lacked an effect on spheroid size even though there was reduced metabolic activity. This at least denotes some potential biological effect of the combined treatment on HER2-positive breast cancer cells. These differing outcomes may be attributed differences in TP53 mutational status, downstream senescence signaling across BC subtypes and differences in crosstalk with primary osteoblasts.[82]

Although this study focused on the senolytics o-Vanillin and RG-7112, there are several other types of senolytic and senomorphic agents acting through distinct pathways. Clinically relevant senolytics such as Navitoclax (ABT-263), Venetoclax (ABT-199), and Dasatinib-based combinations have previously demonstrated efficacy in targeting senescent cancer cell populations and enhancing chemotherapeutic responses in multiple preclinical models.[83–88] Nonetheless, our previous work combining o-Vanillin and RG-7112 with DOX in triple-negative BC co-cultures,[34] together with complementary studies using these senoltyics in musculoskeletal degeneration,[39, 40] further supports the notion that the o-Vanillin/RG-7112 combination can effectively and safely eliminate therapy-induced or age-associated senescent cells and reduce SASP burden to improve treatment efficacy. Our findings position o-Vanillin and RG-7112 as an effective combination that engages both p53-dependent senescent cell clearance (particularly in the stromal compartment) and downregulates several SASP factors associated with treatment resistance and disease progression. Furthermore, our spheroid co-culture model shows good potential for representing a bone-like microenvironment for screening therapeutics against multiple BC subtypes.

Our study marks an important step towards potential translation of combining RG-7112 and o-Vanillin as adjuvant to standard chemotherapy such as DOX. However, an important aspect of these drugs must be noted. RG-7112 is a member of the p53/MDM2 complex inhibitors, derived from the nutlin family of cis-imidazoline analogs designed as anti-cancer agents. RG-7112 underwent clinical trials for leukemia [44], but it lacked specificity and ∼50mg/kg doses and schedule led to several adverse events, preventing it from passing FDA approval [45, 89]. However, our group has shown that 10 times less RG-7112 on a reduced schedule can safely target senescent cells *in vitro*, *ex vivo and in vivo,* without toxicity to non-senescent cells [56, 90]. Natural flavonoids such as o-Vanillin and curcumin have both senolytic and anti-inflammatory properties [91], and they have been tested for potential efficacy in a wide range of disorders with an inflammatory component, including cancers [92–96]. We found that o-vanillin can also kill senescent cells in musculoskeletal tissues [97]. By using the lower, senolytic concentrations of these drugs, we showed that removal of senescent stromal and cancer cells can alter several SASP and inflammatory factors in our breast-to-bone meatastatic microenvironment co-cultures. Future work will assess the impact of this combination on disrupting tumor maintenance and recurrence in the metastatic niche i*n vivo*.

While useful as a platform for evaluating senolytic combination treatments, the 3D co-culture system also has some inherent limitations. The system does not capture systemic features such as immune cell interactions, vascularization, blood flow, or pharmacokinetics, and relies on established cell lines rather than patient-derived tumor cells, thus limiting representation of interpatient heterogeneity. In addition, SASP profiling was restricted to selected factors and may not encompass the full complexity of senescence-associated signaling. Future studies will therefore need to incorporate *in vivo* models of BC bone metastasis, immune-competent systems, and expanded senolytic repertoires to evaluate effects on tumor progression, angiogenesis, immune modulation, and metastatic dissemination. Taken together, this work suggests that senolytic-combination strategies may be a promising potential adjuvant to standard chemotherapy to improve breast-to-bone metastases patient outcomes.

In conclusion, this work demonstrates that TIS arises within both tumor and stromal compartments in a bone-relevant metastatic breast cancer context and that senolytic intervention can mitigate its deleterious consequences of senescence and SASP. By integrating molecularly distinct breast cancer subtypes with primary human spine osteoblasts, this study advances a human-relevant 3D platform for interrogating senescence dynamics within the bone metastatic niche and indicate potential positive targeting of triple-negative and hormone responsive BC in this niche. Our findings provide mechanistic and functional evidence that targeting senescent cells and their associated secretory factors can enhance chemotherapeutic response and suppress pro-tumorigenic signaling. More broadly, this work supports senescence-targeting strategies as a promising avenue for improving therapeutic durability in metastatic breast cancer, and may be clinically applicable to disrupting metastatic site maintenance following chemotherapy. The work presented here establishes a foundation for future *in vivo* and translational studies aimed at limiting treatment-associated disease progression, therapy resistance and relapse.

## Acknowledgements

This project was funded by Cancer Research Society/Canadian Institutes of Health Research operating grant ID #944456, awarded to MHW, LG, LH and DHR. We are thankful for support from Transplant Quebec for patient specimens. DHR gives special thanks to continued support from the McGill Scoliosis and Spine Group. ECBH received and is thankful for the fellowships from the Research Institute of McGill University Health Centre.

## Funding

Funding for this work was obtained from The Cancer Research Society and CIHR 2022 joint operating grant awarded to DHR, LH, MHW and LG.

## Data

The data generated in this study are available upon request from the corresponding authors.

## Conflicts

The authors declare no conflicts of interest.

## Supplementary Materials

**Figure S1.**
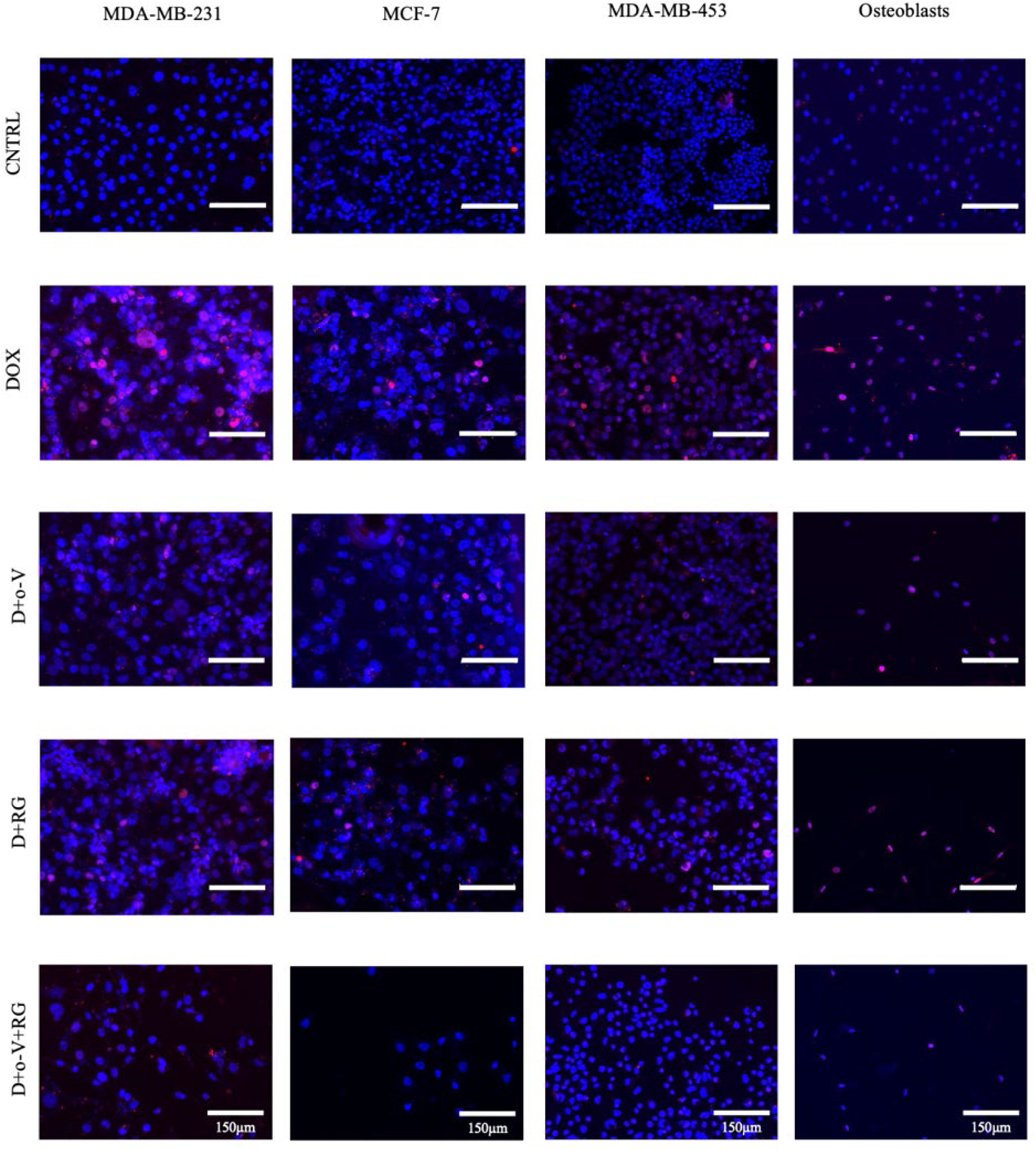
DOX induction and senolytic blockage of senescence in treated breast cancer subtypes and primary spine osteoblast cells in 2D. Representative photomicrographs of p21 (red nuclear staining) expression in cells. DAPI staining was used for nucleus localization. Treated with 0.25μM Doxorubicin alone, or in combination with 5μM RG-7112, and/or 100μM o-Vanillin for 72h.

**Figure S2.**
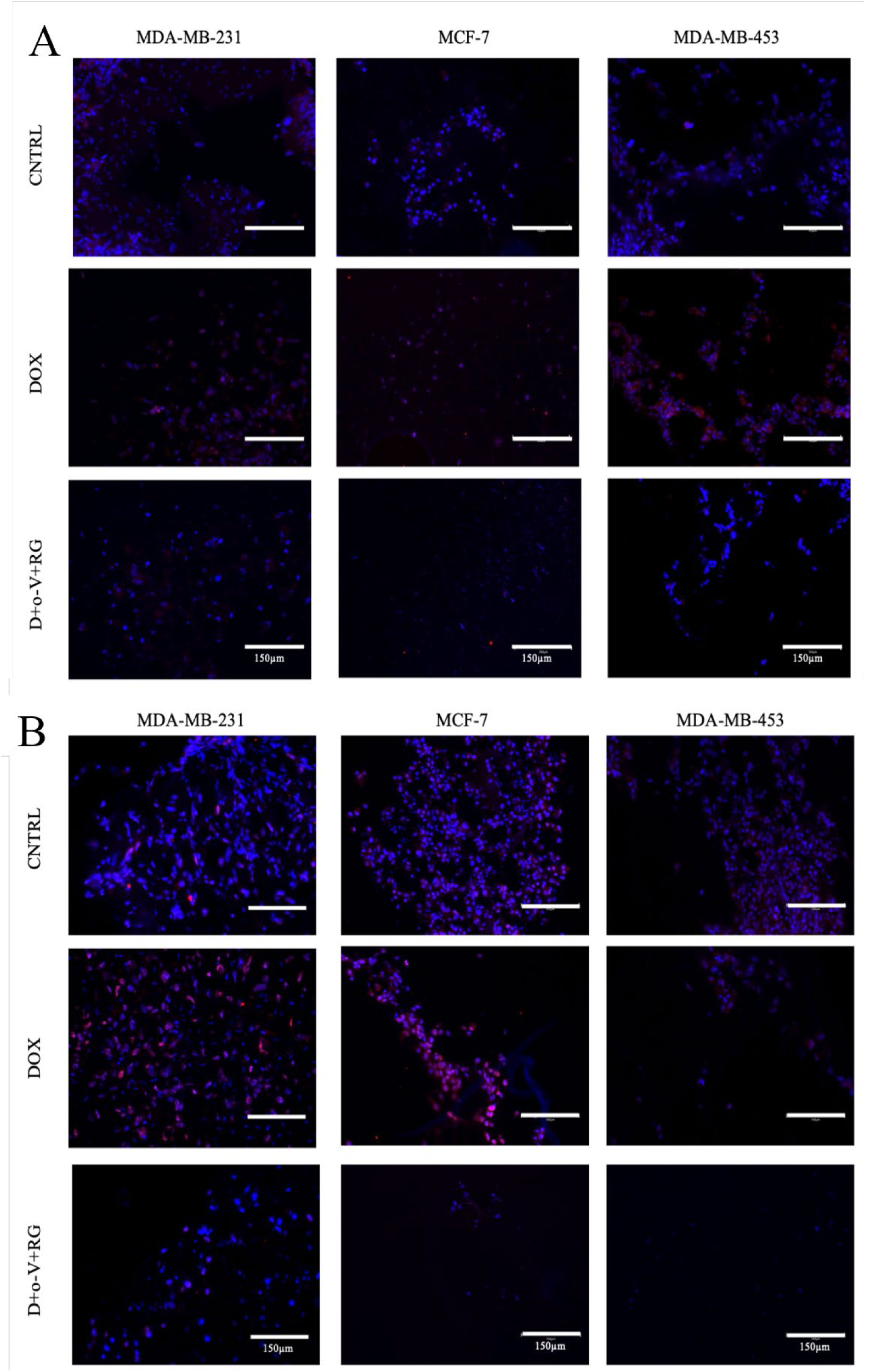
DOX induction and senolytic blockage of senescence in treated breast cancer subtypes and osteoblast 3D co-culture. Representative photomicrographs of A) p21and B) p53 immunofluorescence staining. Treated with 0.5μM Doxorubicin alone or in combination with 5μM RG-7112, and 100μM o-Vanillin for 14days.

## Notes

### Competing Interest Statement

The authors have declared no competing interest.

